# Gradient-aware modeling advances AI-driven prediction of genetic perturbation effects

**DOI:** 10.1101/2025.10.03.680360

**Authors:** Dixian Zhu, Livnat Jerby

## Abstract

Predicting the transcriptional effects of genetic perturbations across diverse contexts is a central challenge in functional genomics. While single-cell perturbational assays such as Perturb-seq have generated valuable datasets, exhaustively profiling all perturbations is infeasible, underscoring the need for predictive models. We present GARM (Gradient Aligned Regression with Multi-decoder), a machine learning (ML) framework that leverages gradient-aware supervision to capture both absolute and relative perturbational effects. Across multiple large-scale datasets, GARM consistently outperforms leading approaches—including GEARS, scGPT, and GenePert—in predicting responses to unseen perturbations within and across contexts. Complementing this, we show that widely used evaluation metrics substantially overestimate performance, allowing trivial models to appear predictive. To address this, we introduce perturbation-ranking criteria (PrtR) that better reflect model utility for experimental design. Finally, we provide insight into gene-specific predictability, revealing pathways and gene classes systematically easier or harder to predict, with implications for model development and biological interpretation. Together, these advances establish a unified methodological and conceptual framework that improves perturbation modeling, sets rigorous evaluation standards, and provides biological insight into gene-specific predictability in functional genomics.

---

Predicting the effects of genetic perturbations and variation is a central challenge in bioscience. Genetic tools, such as CRISPR, allow precise gene knockout, repression, or activation at scale^1–3^, and can be applied in a pooled manner to simultaneously assay thousands of perturbations for their impact on cellular phenotypes. In a Perturb-seq setting, pooled CRISPR screens are performed with single-cell genomics readouts, capturing both the sgRNA and transcriptome (or other molecular modalities) within each cell^4–8^. Yet, perturbational effects are context-dependent, and, despite these advances, exhaustively profiling all genetic perturbations across diverse cell types and conditions is infeasible, underscoring the need for predictive models.

Perturb-seq and other types of perturbational gene expression data have been used to train ML models to predict perturbation effects, with methods as CPA (compositional perturbation autoencoder)^9^ and scGen (single-cell generative model)^10^. More recently, methods such as GEARS (gene activation and repression simulator)^11^, scGPT (single-cell generative pre-trained transformer)^12^, and GenePert (gene perturbation model)^13^ have been developed, with a greater emphasis on unseen genetic perturbations. GEARS, scGPT, and GenePert leverage prior information (e.g., Gene Ontology feature, gene feature pre-trained from a large scale of RNA-Seq data, gene text feature from ChatGPT) to model gene functions and further train the model on Perturb-seq datasets to predict the effect of perturbing these genes.

All of these methods formulate the prediction task as a regression task, training the model to predict the delta in gene expression observed in the perturbed versus (vs.) control cells to be as close as possible to the observed delta based on pointwise regression loss, such as mean squared error (MSE), mean absolute error (MAE), or other variants (**Fig. 1a)**. We have recently shown that this definition of loss is suboptimal in capturing pairwise relationships across datapoints, such that explicitly training predictive models on these relationships (which can be viewed as the “shape” of the data) results in better performances with the same linear time computational complexity as pointwise loss^14^ (**Fig. 1b-c**).

**Figure 1.**
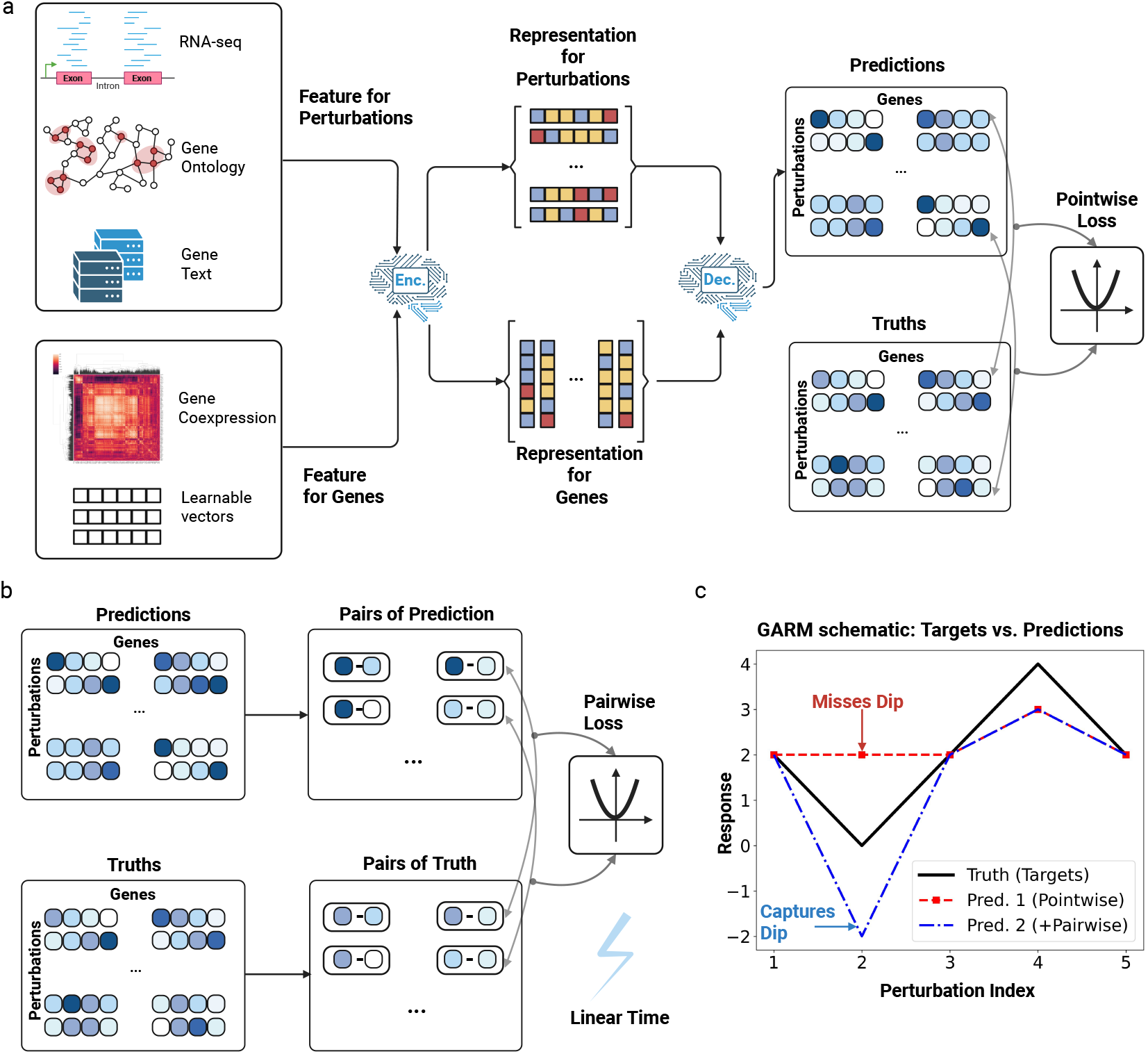
Pairwise loss uncovers perturbational structure overlooked by pointwise regression. **(a)** General learning paradigm for Perturb-seq prediction task. Features for perturbations and genes are taken as input, processed by an encoder module, which learns latent representations. The latent representations are fed to a decoder module (e.g., matrix multiplication or multi-layer perceptron) to predict the perturbation by gene matrix. Previous methods utilize pointwise regression loss, such as MSE, to optimize the model. **(b)** The pairs of prediction differences and measurement differences are constructed as the inputs for the pairwise loss function, where all pairs are used by default with linear time cost. **(c)** Example demonstrating the value of pairwise loss and GARM. Ground truth responses are shown as a solid black line. Prediction #1 (pointwise, red dashed) fails to capture the dip at x=2, while Prediction #2 (pairwise, blue dashed) preserves this relative structure. Prediction 1 and 2 have equal pointwise error but prediction 2 has smaller pairwise error. Pairwise supervision enables models to recover perturbation-to-perturbation relationships critical for PrtR performance.

Model evaluation has likewise relied on metrics of absolute error (e.g., MSE) and metrics that consider the ranking of genes per perturbation. Although widely used, the gene-ranking (GenR) criterion can misleadingly reward trivial models—for example, those predicting the same response for all perturbations—despite offering little value for experiment design or functional inference. For experimental applications, a more relevant question is: *which perturbations are most likely to elicit a desired response in a gene (or gene set) of interest?* To address this, we define perturbation-ranking (PrtR), which evaluates a model’s ability to rank perturbations by their impact on individual genes, complementing the standard absolute value and GenR metric.

Here, we introduce GARM (Gradient-aligned Regression with Multi-decoder), a framework that incorporates gradient-aware supervision and optimizes both absolute and relative prediction objectives. With focus on perturbation modeling we show that gradient-aligned supervision provides a broadly applicable approach for improving AI models whenever learning the relationships between data points is as important as predicting their individual outcomes. Introducing PrtR for model evaluation, we (1) show that GARM consistently outperforms existing approaches (GEARS, scGPT, GenePert, and Coexpress) across multiple Perturb-seq datasets and (2) reveal systematic variation in gene-level predictability across all methods, pointing to biological pathways and gene classes that are intrinsically easier or harder to model. Together, these advances establish a methodological foundation for more powerful predictors, more rigorous evaluation standards, and robust AI-driven experimental design.

## RESULTS

### Gradient Aligned Regression with Multi-decoder (GARM)

Prior ML models for perturbational data prediction were optimized to minimize pointwise regression loss, such as MSE or MAE. To better capture the pairwise relationship between different perturbations, we propose to utilize pairwise regression losses to predict Perturb-seq data (**Fig. 1b**). We have previously shown analytically that training deep learning models with pairwise losses is equivalent to learning the gradient of the ground truth function and beneficial for capturing the relationship between datapoints^14^. As a practical example, the pairwise losses prefer the ‘shape-consistent’ predictions that maintain the ranks from the ground truths (**Fig. 1c, Extended Data Fig. 1c**).

The general format for pairwise loss is defined as:

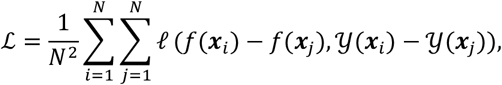

where ℓ (·,·) is a standard loss function applied to the prediction f (·) and measured data 𝒴 (·) when considering a pair of perturbations *x*_*i*_ and *x*_*j*_.

For predicting the impact of *P* genetic perturbation on the expression of *G* genes, we consider all pairs of genes per perturbation (*PG*^*2*^) and all pairs of perturbations per gene (*GP*^*2*^) when calculating pairwise losses, to learn the pairwise relationships between different pairs of genes and different pairs of perturbations. By leveraging the equivalent forms of the objective functions for the pairwise losses^14^, the quadratic computational cost part can be accelerated to linear cost, i.e. *O*(*PG*^2^ + *GP*^2^) → *O*(*GP*). The user can also define which types or subsets of pairs to include as an input, providing flexibility in cases where there are certain aspects that are more important to capture (e.g., focus only on perturbation ranking per gene).

Here, we further developed our previous implementation of these principles (GAR^14^) into a more efficient pairwise regression method, named GARM (**Fig. 2a, Extended Data Fig. 1a-b)**. GARM does not require method-specific hyper-parameter tuning, and still maintains or surpasses the performances obtained with method-specific hyper-parameter tuning (**Extended Data Fig. 2, 3a**). GARM is based on a multi-decoder structure that generates and consolidates multiple versions of the prediction (**Fig. 2a**). Five different decoders are trained with different data arrangements and loss functions (**Extended Data Fig. 1a, Methods**), focused on a different aspect of the Perturb-seq prediction task, that is, minimizing the MAE (1), optimizing PrtR (2-3), and optimizing GenR (4-5). The five different predictions are then reconciled to generate the final output (**Extended Data Fig. 1b, Methods**). As described below, GARM was tested on two synthetic datasets (**Extended Data Fig. 2**) and four representative Perturb-seq datasets in comparison to four other methods in predicting perturbational effects in the same and unseen cellular contexts.

**Figure 2.**
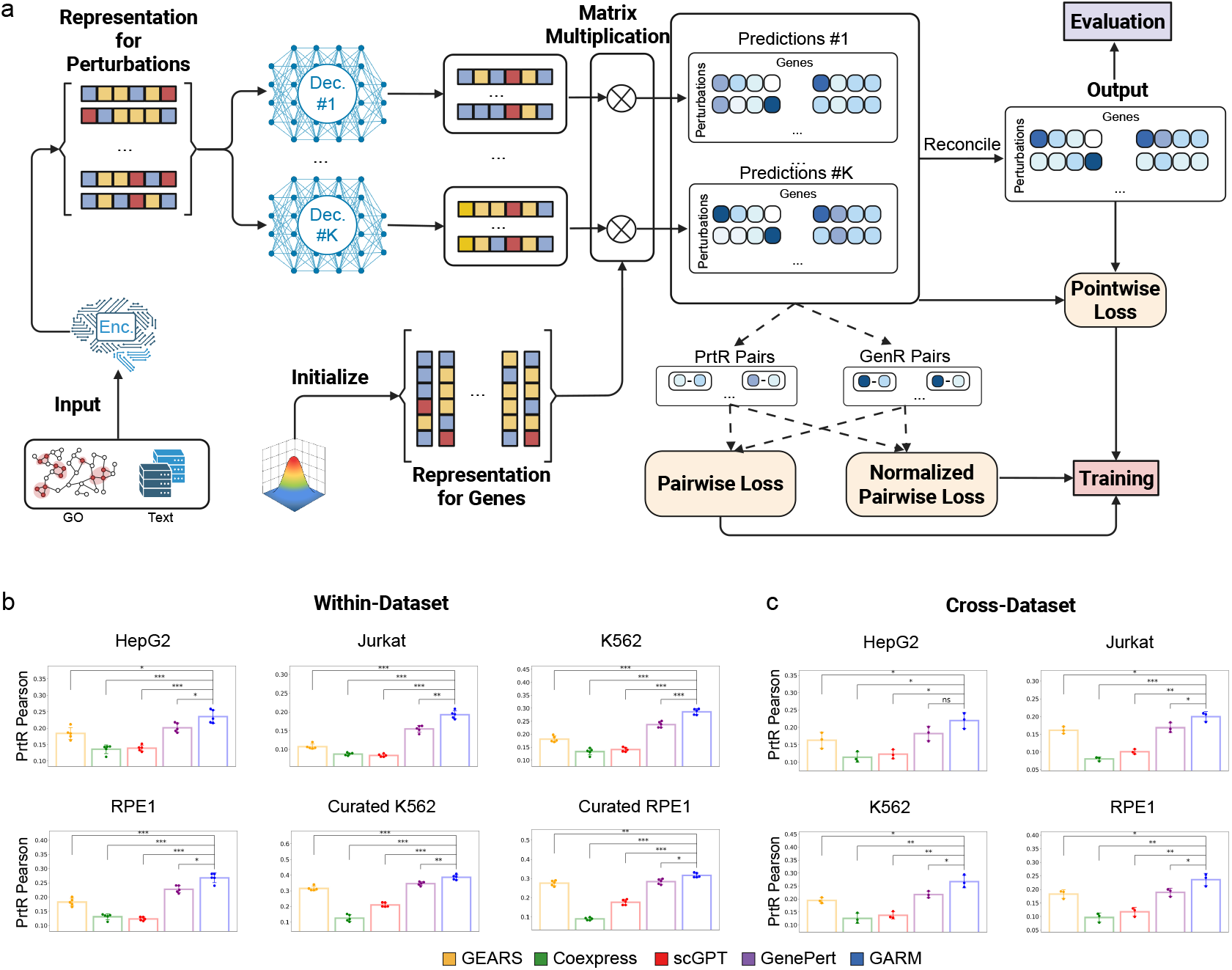
GARM outperforms state-of-the-art predictors across Perturb-seq datasets. **(a)** GARM overview: like previous methods, the encoder takes GO and text data as input; multiple decoders generate predictions based on the GAR pairwise loss minimization; using a multi-decoder design, multiple decoders are trained simultaneously, and the predictions are reconciled to generate the final output, not relying on any hyperparameter to balance the different objectives. (**b)** Comparison of different methods in the Within-Dataset setting for PrtR Pearson correlation coefficients evaluation (mean ± standard deviation (SD)), showing the 5 performance values from the 5 data splits on different random seeds. **(c)** Comparison of different methods in the Cross-Dataset setting for PrtR Pearson correlation coefficients evaluation (mean ± SD), showing the 3 performance values from the 3 data splits on the different choices for training and validation. For all the barplots, ^***^*P* < 0.0001, ^**^*P* < 0.001, ^*^*P* < 0.05, one-sided t-test.

### Predicting transcriptional response to unseen genetic perturbations within the same dataset

GARM was benchmarked against four other methods developed to predict Perturb-seq data: GEARS^11^, scGPT^12^, GenePert^13^, a coexpression based linear model (termed as ‘Coexpress’ for simplicity)^15^. Each method is tuned with its hyper-parameters and run independently with five different random seeds (**Methods**). To assess evaluation metrics, a model that predicts all perturbations to have the same impact on the cell transcriptome was also applied, such that the average transcriptional response observed across all the perturbed cells in the training data is used to predict the transcriptional response to any perturbation.

Four large-scale CRISPRi Perturb-seq datasets were used: HepG2^7^, Jurkat^7^, K562^6^, RPE1^6^, each spanning 2,042 - 2,373 perturbations, with the latter two also used in their curated version, where only 1,087 – 1,534 strong perturbations and the 5,000 topmost variable genes were retained^11^. Performances were evaluated based on the standard MSE and GenR, extensively used in prior studies, and PrtR, proposed here (**Table 1**).

**Table 1.**
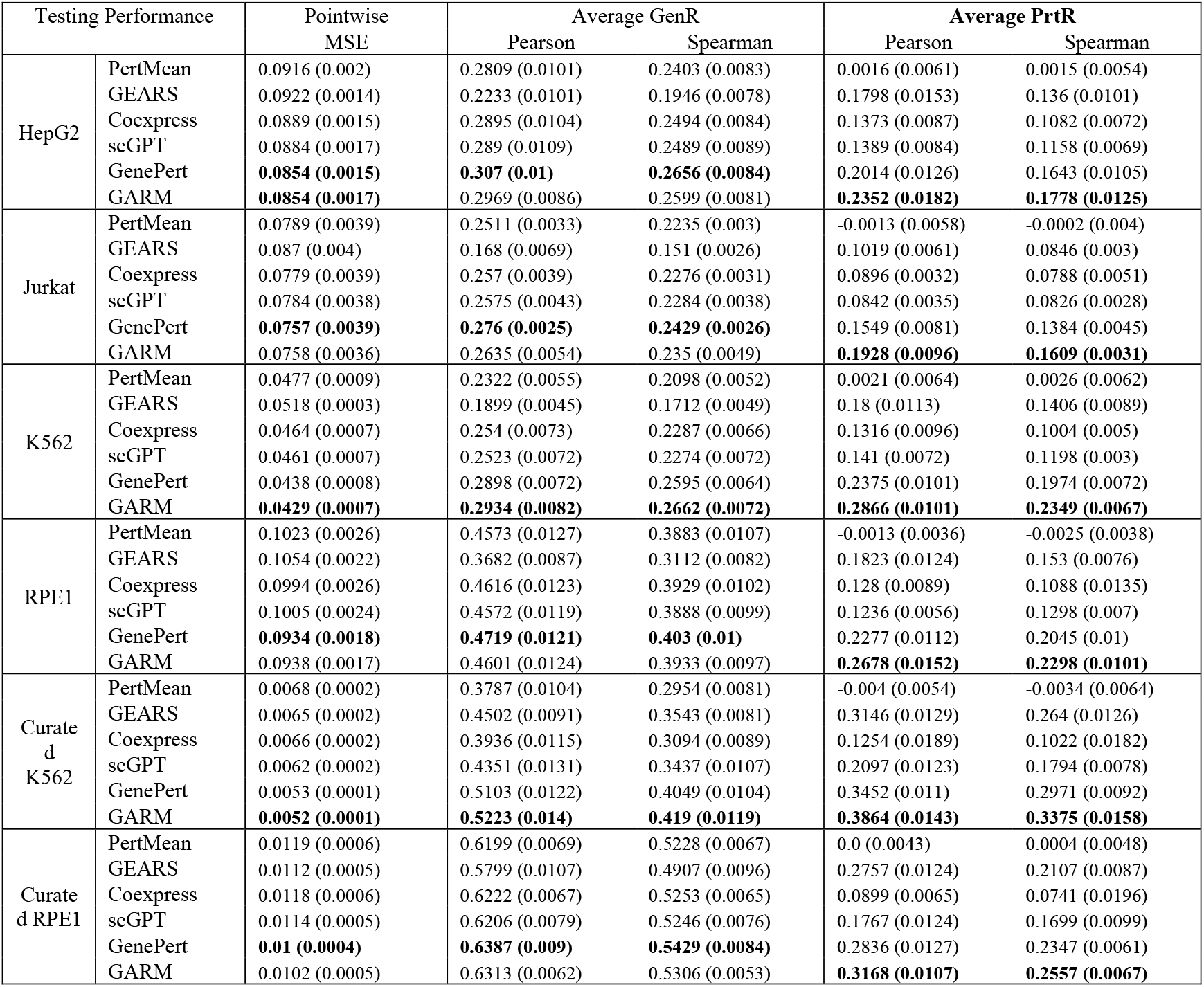
Method performances in the within-dataset setting. The reported values are in mean (standard deviation) format across the 5 averaged values from the 5 different random splits of the data.

GARM significantly outperforms all other methods when considering the average PrtR Pearson correlation coefficients in all 6 datasets and in 5 out of the 6 datasets based on average PrtR Spearman (**Table 1, Fig. 2b, Extended Data Fig. 3b**). Evaluating PrtR model performances per gene, shows that GARM outperforms all other methods in the majority of genes, both based on Pearson (**Fig. 3a,c**) and Spearman correlation tests (**Fig. 3c, Extended Data Fig. 3c**). More specifically, GARM outperformed all other competitors in the PrtR task with Pearson correlation in 84.5%, 87.0%, 90.6%, 89.9%, 86.5%, 72.5% of genes and with Spearman correlation in 66.3%, 80.9%, 82.9%, 83.9%, 84.5%, 69.1% of the genes when tested on HepG2, Jurkat, K562, RPE1, Curated K562, and Curated RPE1 datasets, respectively. GARM also maintains high performance based on MSE and GenR, being the best or among the top two methods in all datasets (**Table 1**).

**Figure 3.**
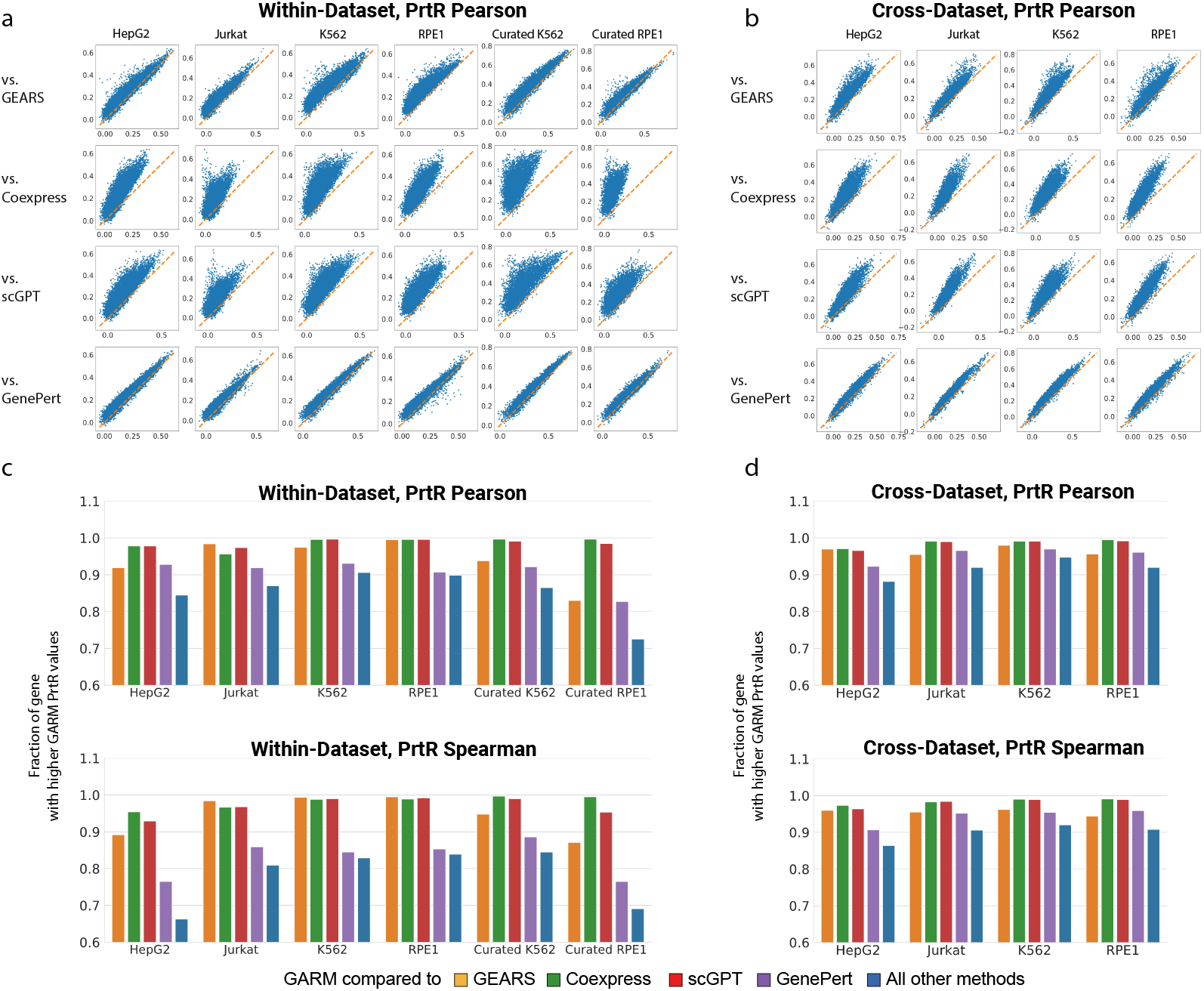
Gene-level evaluation highlights GARM’s predictive advantage and broad variability in predictability across genes. **(a-b)** PrtR Pearson correlation coefficients performances comparison for GARM vs. other competitors at the gene level: Each dot corresponds to a gene, showing GARM PrtR performance (y axis) and the alternative method PrtR performance (x axis). **(a)** Within-Dataset setting, showing the average performance value obtained per gene across 5 different random data splits per dataset. **(b)** Cross-Dataset setting, showing the average performance value across the 3 data splits. **(c-d)** Fractions of genes for which GARM outperformed the other method based on the PrtR metric in within-dataset **(c)** and cross-dataset **(d)** settings, when quantifying PrtR based on Pearson (top) and Spearman (bottom) correlation coefficients.

This also shows that PrtR is a more difficult task, with performances for unseen perturbation prediction within-dataset ranging from 0.0842 to 0.3864 across all datasets and five methods (GARM, GEARS, scGPT, GenePert, and Coexpress) for average Pearson correlation coefficients, as opposed to 0.168 to 0.6387 in the GenR task (**Table 1**). Moreover, a model that predicts the mean value of all training perturbations as the perturbational effect for all perturbations is successful in the GenR task and even outperforms all other methods in certain settings (**Table 2**).

**Table 2.** Method performances in cross-dataset setting. The reported values are in mean (standard deviation) format across the 3 averaged values from the 3 training-validation data splits.

| Testing Performance |  | Pointwise MSE | Average GenR |  | Average PrtR |  |
| --- | --- | --- | --- | --- | --- | --- |
|  |  |  | Pearson | Spearman | Pearson | Spearman |
| HepG2 | PertMean | 0.1017 (0.0008) | 0.2207 (0.026) | 0.1839 (0.0168) | -0.0003 (0.0031) | -0.0011 (0.0026) |
|  | GEARS | 0.1059 (0.0019) | 0.2298 (0.0316) | 0.1806 (0.0272) | 0.1629 (0.0189) | 0.1411 (0.0194) |
|  | Coexpress | 0.1009 (0.0005) | 0.2283 (0.0264) | 0.192 (0.0182) | 0.1142 (0.0122) | 0.0914 (0.0087) |
|  | scGPT | 0.1014 (0.0021) | 0.2288 (0.024) | 0.1954 (0.0178) | 0.1227 (0.0106) | 0.1105 (0.0094) |
|  | GenePert | 0.1002 (0.0002) | 0.2367 (0.0279) | 0.2017 (0.0214) | 0.1823 (0.0168) | 0.1574 (0.0164) |
|  | GARM | **0.0959 (0.0026)** | **0.2505 (0.0334)** | **0.2155 (0.0252)** | **0.2198 (0.0183)** | **0.1929 (0.0187)** |
| Jurkat | PertMean | 0.0942 (0.0034) | 0.1629 (0.0104) | 0.1561 (0.0075) | 0.0011 (0.0029) | 0.0005 (0.0028) |
|  | GEARS | 0.0997 (0.0056) | 0.1902 (0.0102) | 0.1599 (0.0048) | 0.1615 (0.0079) | 0.1403 (0.0097) |
|  | Coexpress | 0.0941 (0.0037) | 0.1715 (0.0096) | 0.1624 (0.0086) | 0.0804 (0.0047) | 0.0728 (0.0041) |
|  | scGPT | **0.091 (0.0014)** | 0.1758 (0.0101) | 0.1655 (0.0078) | 0.1005 (0.0057) | 0.0889 (0.0042) |
|  | GenePert | 0.0912 (0.0022) | 0.1973 (0.0077) | 0.1726 (0.0077) | 0.1684 (0.0111) | 0.1476 (0.011) |
|  | GARM | 0.0931 (0.005) | **0.199 (0.0103)** | **0.1808 (0.0082)** | **0.1999 (0.0111)** | **0.1852 (0.013)** |
| K562 | PertMean | 0.0604 (0.0041) | 0.1465 (0.0114) | 0.1417 (0.0082) | -0.0028 (0.0044) | -0.0022 (0.004) |
|  | GEARS | 0.0699 (0.0062) | 0.1935 (0.0073) | 0.1574 (0.0031) | 0.1943 (0.0088) | 0.1637 (0.0091) |
|  | Coexpress | 0.0607 (0.0044) | 0.1605 (0.0094) | 0.1547 (0.0066) | 0.1256 (0.0156) | 0.1031 (0.013) |
|  | scGPT | **0.0555 (0.0028)** | 0.1682 (0.0095) | 0.1606 (0.0069) | 0.1372 (0.0111) | 0.1199 (0.0088) |
|  | GenePert | 0.0561 (0.0034) | 0.1922 (0.0107) | 0.1726 (0.0076) | 0.2173 (0.0105) | 0.1809 (0.0101) |
|  | GARM | 0.0598 (0.0048) | **0.199 (0.0112)** | **0.1964 (0.0084)** | **0.2671 (0.02)** | **0.2247 (0.0151)** |
| RPE1 | PertMean | 0.1247 (0.0031) | **0.3378 (0.0439)** | **0.281 (0.0271)** | 0.0006 (0.0014) | 0.0006 (0.0014) |
|  | GEARS | 0.1226 (0.0053) | 0.2734 (0.03) | 0.2164 (0.0225) | 0.1828 (0.0129) | 0.1595 (0.0128) |
|  | Coexpress | 0.1221 (0.004) | 0.3235 (0.0404) | 0.2768 (0.0275) | 0.0966 (0.0135) | 0.0844 (0.0115) |
|  | scGPT | 0.1252 (0.0026) | 0.3233 (0.0315) | 0.2768 (0.0246) | 0.1174 (0.0132) | 0.1078 (0.01) |
|  | GenePert | 0.1208 (0.004) | 0.2975 (0.0251) | 0.2723 (0.0254) | 0.1892 (0.0117) | 0.1648 (0.0121) |
|  | GARM | **0.1177 (0.0053)** | 0.2979 (0.0327) | 0.2554 (0.024) | **0.236 (0.0184)** | **0.2118 (0.0174)** |

For experimental design, it is often useful to identify a subset of perturbations that up- or down-regulate a gene of interest for subsequent testing, while the specific ranking of perturbations that do not have a significant effect on the gene is less informative. Thus, predictors were also tested by defining, per gene, the top 20% of perturbations that up-regulate it and the top 20% of perturbations that down-regulate it, and examining whether the predicted values correctly identify these perturbations by computing the Area Under the Receiver Operating Characteristic Curve (AUROC) and Area Under the Precision Recall Curve (AUPRC, **Supplementary Table 1**). GARM outperformed all other methods when tested based on the AUROC and AUPRC evaluation criteria under all 24 PrtR evaluations.

### Predicting transcriptional responses to genetic perturbations in unseen cellular contexts

As Perturb-seq screens can be performed genome-wide in a single or a small number of cellular contexts and conditions, another important task is to use such data to predict the perturbational effects in a new, unseen context. GARM outperformed other methods also in such settings.

For cross-dataset prediction evaluation, four large essential-wide CRISPRi Perturb-seq datasets were used, namely, HepG2^7^, Jurkat^7^, K562^6^, and RPE1^6^, focusing on 6641 genes included in all processed datasets. In each iteration, one of the four datasets was used for testing, one for validation, and the remaining two datasets for training, resulting in 12 different data splits. For each dataset used as the testing set, the performance was evaluated based on the three different training and validation data splits.

As in the within-dataset setting, GARM was compared to GEARS^11^, scGPT^12^, GenePert^13^, and Coexpress^15^, and performances were evaluated based on the standard MSE, GenR, and PrtR (**Table 2**). GARM outperforms all other methods in the PrtR task (**Table 2**), with significant improvement compared to all other methods in 3 out of the 4 datasets (on both Pearson and Spearman correlations, **Fig. 2c, Extended Data Fig. 3d**). GARM maintains high performance also based on the MSE and GenR tasks (best based on GenR in 3 out of the 4 datasets, **Table 2**). At the single gene level, GARM outperformed all methods in ranking the perturbation effects on 88.2%, 92.0%, 94.8%, 92.0% of the genes based on Pearson correlation tests (**Fig. 3b,d**) and 86.4%, 90.6%, 92.0%, 90.8% of the genes based on Spearman correlation tests (**Fig. 3d, Extended Data Fig. 3e**) in the HepG2, Jurkat, K562, and RPE1 Perturb-seq datasets, respectively. GARM consistently outperformed all methods under the AUROC and AUPRC evaluation criteria defined before in predicting each one of the four datasets (**Supplementary Table 2**).

### The predictability of the transcriptional response to a genetic perturbation varies substantially across genes

The variation in prediction performances does not only vary by the model used, but also by gene, in a manner that is consistent across models and datasets (**Fig. 3a-b, Extended Data Fig. 3c,e**). Certain genes show low PrtR values across all methods and datasets (i.e., are harder to predict), while others show high PrtR values across all methods and datasets (i.e., are easier to predict). GARM obtained more accurate predictions for the majority of genes (i.e., higher PrtR values) compared to all other methods (**Fig. 3c-d**), such that the GARM PrtR value of a gene is correlated with its PrtR value obtained with other methods (**Fig. 3a-b**).

This raises the question: Why are some genes harder to predict than others? Gene set enrichment analyses (GSEA) on the PrtR values obtained in the within-dataset setting for GARM show that genes with high PrtR values are significantly enriched for genes involved in DNA packaging, cell cycle, apoptosis, metabolism, antigen presentation, and other processes (**Methods, Fig. 4a, Supplementary Table 3**). Genes with low PrtR values show a weaker connection to gene set annotations, but are enriched for transcription factors, chromatin modifiers, and epigenetic regulators (**Methods, Supplementary Table 3**). However, more than half of the gene sets show a variation in PrtR correlation values, spanning the full spectrum of low (< 0.1) and high (> 0.4) values, with no significant skew toward higher or lower PrtR values, indicating that this property depends on additional factors.

**Figure 4.**
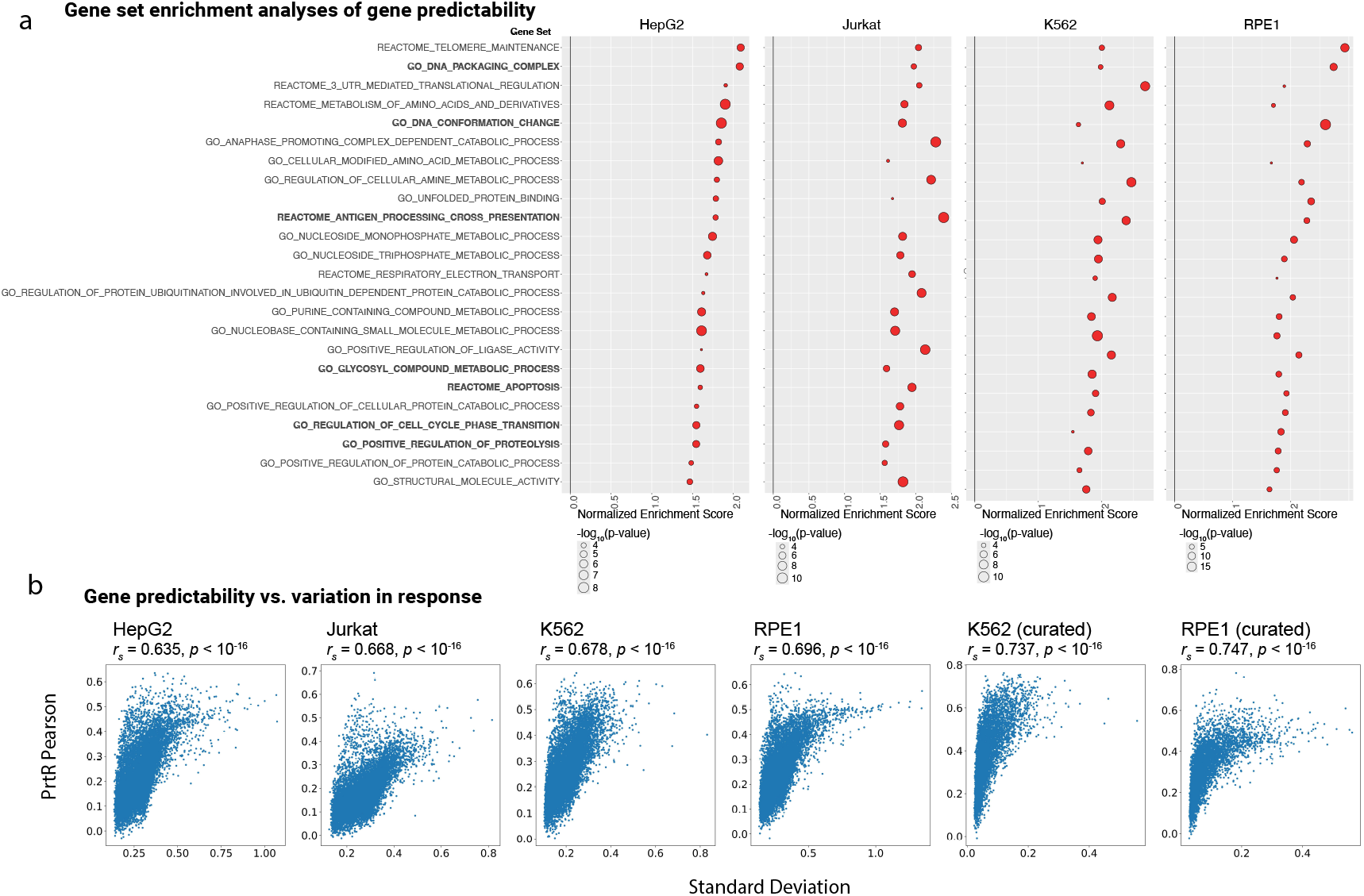
Gene-level predictability reveals pathways systematically easier—or harder—to model. **(a)** Predictable genes are enriched for specific gene sets: GSEA was performed for each of the 4 datasets (HepG2, Jurkat, K562, and RPE1) to identify gene sets showing significantly higher or lower than expected GARM PrtR Pearson correlation coefficients. Gene sets showing a significant enrichment based on all 4 datasets are shown (y axis) with the matched normalized enrichment score (x axis) and p-value (dot size). **(b)** Gene predictability is associated with the magnitude of the gene responsiveness to genetic perturbations: GARM PrtR Pearson correlation coefficients (y axis) versus standard deviation (std) of each gene in the Perturb-seq data (pseudo-bulk log1p-transformed tpm; x axis), shown per gene (dot) in each of the datasets. The Spearman correlation between the two values in each dataset is noted.

Genes whose expression is more substantially impacted by genetic perturbations (quantified as the standard deviation of the gene in response to all perturbations in a given dataset, **Methods**) are more likely to have higher PrtR values (**Fig. 4b**). While this may be beneficial in the sense that the model is capturing the strong signals in the data and does not fit to changes that may be more prone to technical noise and stochasticity, this may also indicate that more subtle (yet important) effects are still not optimally modeled and predicted by existing models. GARM improves the prediction accuracy also for these genes, yet it may be necessary to provide the model with additional types of prior data to reach greater predictability (e.g., PrtR correlation coefficient > 0.4) for this class of genes.

## DISCUSSION

The work presented here demonstrates the advantages of directly incorporating pairwise relationships into predictive model training and optimization. GARM operationalizes this principle through gradient-aligned supervision and a multi-decoder design, consistently outperforming four state-of-the-art methods in predicting cellular transcriptional responses to genetic perturbation.

Our study also exposes limitations of evaluation criteria that have been widely used to benchmark predictors of Perturb-seq data^11–13,15^. These criteria overestimate model performances and, as we show via the PertMean model (e.g., **Table 2**, RPE1 GenR values), fail to distinguish between models that capture the impact of the genetic perturbation per se and models that simply overfit to general trends that are not perturbation-specific and more likely result from technical noise and stochastic variation. By contrast, the perturbation-ranking (PrtR) evaluation introduced here provides a more stringent and biologically relevant benchmark.

From a forward genetics perspective, the goal is often to identify perturbations that alter the cell transcriptome in a specific manner, for example, leading to specific cell state transitions of interest. Yet Perturb-seq screens are typically limited by the number of perturbations that can be tested. A critical question therefore becomes: *which perturbations should be prioritized to drive specific cell state transitions?* The PrtR criterion explicitly evaluates this ability, making it directly useful for forward-genetics applications and AI-driven design of Perturb-seq experiments.

While prior reports suggested that simple linear models can perform as well as more complex non-linear ones in predicting perturbational effects^15^, GARM demonstrates clear improvements over the Coexpress linear baseline under PrtR evaluation. This result highlights both the added value of non-linear architectures and the importance of using evaluation metrics that more faithfully reflect the difficulty and diversity of the prediction task.

GARM outperforms other methods in most settings and improves the prediction for the majority of genes (**Fig. 3**). However, it is not orthogonal to other methods: genes that are difficult for existing models to predict remain relatively more challenging for GARM. Hard-to-predict genes may highlight biological contexts where the assumptions or priors shared across methods (e.g., coexpression structure, gene set annotations, or text-based embeddings) are insufficient. New approaches or different types of data may be needed to robustly predict the response of such genes to genetic perturbations. Another possibility is that such genes suffer from lower signal-to-noise ratio in Perturb-seq data, owing to dropouts, batch effects, or other confounders. In this sense, improvements in experimental protocols and preprocessing pipelines will be just as critical as methodological advances in improving prediction accuracy.

Beyond its empirical performance, a central conceptual novelty of GARM lies in explicitly aligning learning objectives with the relational structure of the data. By training models not only to minimize pointwise error but also to preserve pairwise differences, GARM captures the geometry of perturbational responses and the relative ordering among perturbations. This focus on relational fidelity distinguishes GARM from prior approaches, enabling it to generalize more effectively to unseen perturbations and contexts. More broadly, incorporating notions of data structure and relativity into loss functions represents a generalizable strategy for predictive modeling in high-dimensional biology.

Many biological questions are inherently comparative—for example, whether perturbation *A* elicits a stronger effect than perturbation *B*, or whether two perturbations drive similar expression programs. By embedding these relational objectives into the loss function, GARM aligns model optimization with the types of inferences researchers seek to make. This ensures that learned representations reflect the geometry of the perturbational landscape, enabling more meaningful generalization to unseen perturbations and supporting tasks such as prioritizing candidates for follow-up experiments or designing minimal yet informative perturbation screens.

Taken together, our study introduces a new conceptual and methodological framework for training and evaluating models of perturbational effects. By integrating gradient-aligned supervision with biologically relevant evaluation, GARM advances the development of predictive models that can accelerate AI-guided experimental design and functional genomics.

## ACKNOWLEDGEMENTS

L.J. is a Chan Zuckerberg Biohub Investigator and an Allen Distinguished Investigator. This study was supported by the Burroughs Wellcome Fund (BWF, 1019508.01; L.J.), National Human Genome Research Institute (NHGRI) IGVF (Impact of Genomic Variation on Function) Consortium (U01HG012069; L.J.), Paul G. Allen Family Foundation (12965; L.J.), and Chan Zuckerberg Biohub (L.J.).

## AUTHOR CONTRIBUTIONS

D.Z. and L.J. conceived and designed the study. D.Z. developed the code and performed the analyses. D.Z. and L.J. interpreted the results and wrote the manuscript. L.J. obtained funding and supervised the study.

## METHODS

### Gradient Aligned Regression with Multi-decoder (GARM)

Let ***y***^*p*^ be the transcriptional response to a perturbation *p*. The prediction task of a predictive Perturb-seq model is to learn a function 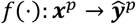, that takes the input data ***x***^*p*^ representing the perturbation *p* (**Fig. 1a**) and predicts the response on gene expressions. *f* is composed of the encoders and decoders.

The standard pointwise regression loss function has been used in previous studies to have the predictions 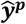 match the ground truth ***y***^***p***^:

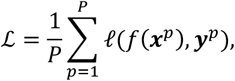

Where *P* is the total number of perturbations and *l* (·,·) is a standard pointwise loss function, such as MAE or MSE loss, i.e. *l*(***a, b***) = ‖***a*** − ***b***‖_1_ or 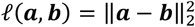.

To better capture pairwise relationship (e.g. perturbation A activate a gene of interest more than perturbation B), GARM incorporates pairwise regression loss functions:

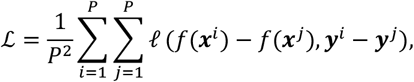

where *l* (·,·) can be defined similarly as pointwise loss. In our previous work^14^, we proved that optimizing the pairwise regression loss equals to optimizing the Pearson correlation coefficients with certain *l*(·,·), which is also related to learning the gradients of the ground truth function.

GARM is designed with five different decoders to generate different versions of prediction, which are optimized with different objectives and then reconciled to produce the final prediction.

#### Prediction #1

Optimized by conventional Mean Absolute Error (MAE) pointwise loss:

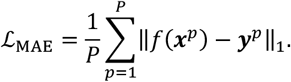

This version of prediction is designed to learn the mean of each target value.

#### Prediction #2

Construct the perturbation ranking (PrtR) pairs per gene and optimize the Mean Squared Error (MSE) based pairwise loss. Deonte the number of expressed gene as *G, g*-th item of prediction as *f*_*g*_, *g*-th item of ground truth as ***y***_*g*_:

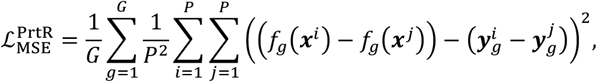

Based on the theorem 1 from the previous work^14^, this pairwise loss is equivalent to:

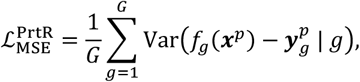

Where Var stands for the empirical variance for the error term, 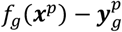, for the random variable ***x***^*p*^ and ***y***^*p*^ conditioned on *g* . It is well-known that computing the variance only requires linear time complexity regarding to data size. Therefore, the total time complexity for computing 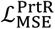 only need O (GP), rather than O (GP^2^).

It is worth noting that 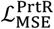 explicitly enforces the prediction #2 captures the pairwise difference for targets on different perturbations, 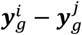 per-gene *g*, which is different with pointwise loss (**Fig. 1c, Extended Data Fig. 1c**). It is also well-known that:

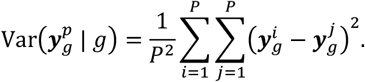

Therefore, compare with pointwise loss, 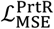 explicitly enforces the prediction to learn and estimate the variance of the target values per-gene better.

#### Prediction #3

Construct the PrtR pairs per gene followed by L2-norm normalization and optimize the L2-norm based pairwise loss. Fixing a gene, *g*, denote the pair of ground truth as 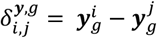, and the pair of the prediction as 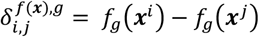. Denote the vector for PrtR pairs of ground truth perturbations on gene *g* as 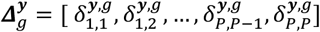, and the vector for PrtR pairs of prediction on gene *g* as 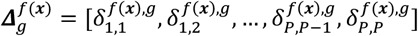. The L2-norm loss on PrtR pairs is defined as following:

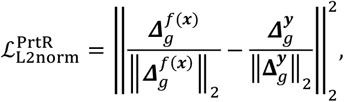

where the L2-norm normalization removes the influence of different magnitudes comparing with 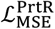.

Follow the Corollary 3 from our previous work^14^, it has the following equivalent form:

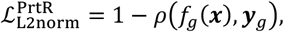

where *ρ*A*f*_*g*_(***x***), ***y***_*g*_B represents Pearson correlation coefficients between the predictions *f*_*g*_(***x***) and the ground truths ***y***_*g*_ on gene *g* for all the perturbations in the training dataset. Therefore, 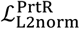 explicitly enforces the prediction to capture the trend of ground truth per-gene better.

#### Prediction #4

Construct the gene ranking (GenR) pairs per perturbation and optimize the Mean Squared Error (MSE) based pairwise loss (similar but orthogonal to prediction #2):

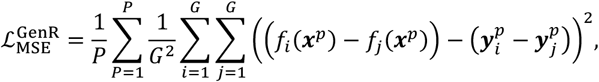

Based on the theorem 1 from the previous work^14^, this pairwise loss is equivalent to:

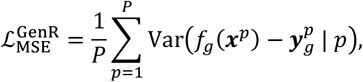

Where Var stands for the empirical variance for the error term, 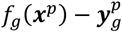, for the random variable *g* conditioned on *p*. It is well-known that computing Var only requires linear time complexity regarding to data size. Therefore, the total time complexity for computing 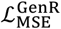 only need O (PG), rather than O (PG^2^).

It is worth noting that 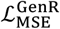 explicitly enforces the prediction #4 captures the target pairwise difference, 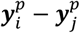 per-perturbation *p*, which is different with pointwise loss (**Fig. 1c, Extended Data Fig. 1c**). It is also well-known that:

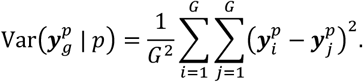

Therefore, compare with pointwise loss, 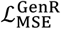 explicitly enforces the prediction to learn and estimate the variance of the target values per-perturbation better.

#### Prediction #5

Construct the GenR pairs per perturbation followed by L2-norm normalization and optimize the L2-norm based pairwise loss. Fixing a perturbation, *p*, denote the pair of ground truth as 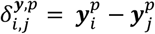, and the pair of the prediction as 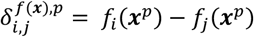. Denote the vector for GenR pairs of ground truth perturbations on perturbation *p* as 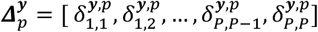, and the vector for GenR pairs of prediction on perturbation *p* as 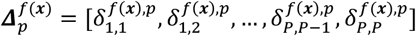. The L2-norm loss on GenR pairs is defined as following:

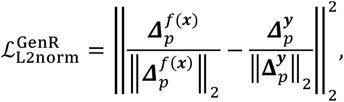

where the L2-norm normalization removes the influence of different magnitudes compared with 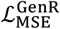.

Follow the Corollary 3 from our previous work^14^, it has the following equivalent form:

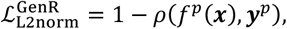

where *ρ*(*f*^*p*^(***x***), ***y***^*p*^) represents Pearson correlation coefficients between the predictions *f*^*p*^(***x***) and the ground truths ***y***^*p*^ on perturbation *p* for all the expressed genes in the training dataset. Therefore, 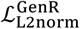 explicitly enforces the prediction to capture the trend of ground truth per-perturbation better. Reconciliations:

The five different versions of predictions capture target mean, target variance per-gene, target trend pergene, target variance per-perturbation, and target trend per-perturbation, and are reconciled as follows. Denote prediction version #1 to #5 as *f* (***x***, #1) to *f*(***x***, #5). Following the conventional notations, denote 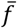 as the mean and *σ*(*f*) as the standard deviation.

#### Step 1: reconcile the variance estimations with the trend estimations

Standardize the prediction #3 per-gene, so that only the trend is kept:

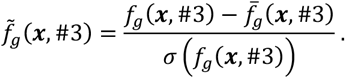

Reconcile the variance from the prediction #2 per-gene with the trend:

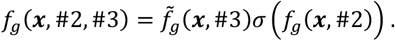

Standardize the prediction #5 per-perturbation, so that only the trend is kept:

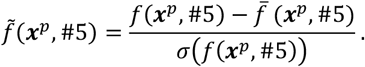

Reconcile the variance from the prediction #4 per-perturbation with the trend:

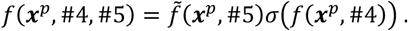

#### Step 2: reconcile the mean estimations with the trend-variance estimations

Calculate and denote the mean estimation per-gene from prediction #1: 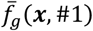. Calculate and denote the mean estimation per-perturbation from the prediction #1: 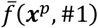. Reconcile the mean from the prediction #1 per-gene with the trend-variance estimations:

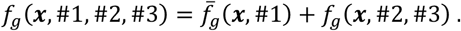

Reconcile the mean from the prediction #1 per-perturbation with the trend-variance estimations:

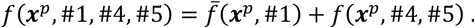

#### Step 3: produce the final prediction with the average of trend-variance-mean estimations from per-gene and per-perturbation processes

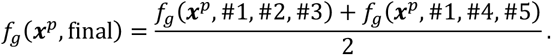

### Testing GARM in comparison to GAR

GARM was benchmarked compared to GAR in fitting to two synthetic datasets (**Extended Data Fig.2**) and four real world Perturb-seq datasets (**Extended Data Fig. 3a**), as described below. Two synthetic datasets were constructed: Sine and Amplified-Sine. The Sine dataset is generated by standard Sine function, *y* = sin *x*, where *x* ∈ [−10π, 10π) with sampling interval as 0.1 and a total of 629 data samples. The Amplified-Sine dataset was generated similarly by a more complicated function, *y* = |*x*| sin *x*, where *x* ∈ [−50, 50) with sampling interval as 0.1 and 1000 data samples. Half of the data was randomly sampled as training, and the remaining half was held out for testing.

The backbone model was chosen as an 8-layered Feed Forward Neural Network (FFNN) with 6 hidden layers, where each hidden layer contains 100 neurons. Exponential linear units (ELU) activation function was applied on each neuron^16^. Each method was run independently five times with different random seeds. The final prediction was averaged across the five independent trials with the standard deviation reported and the evaluation metrics (MAE and Pearson correlation coefficients) computed based on the average prediction (**Extended data Fig. 2**). For each trial included 1200 epochs of training by Adam optimizer^17^ with initial learning rate tuned and set as 1e-3 and decreased by 10 folds at the end of {300, 600, 900}-th epoch. The batch size for each mini-batch iteration was 64. The weight decay was set as 0.

GARM was compared to three variants of GAR (applied with {0.1, 1, 10} as the hyper-parameter α, as done in the original paper^14^), MAE and MSE loss (which are the most typical pointwise regression loss functions), and a heuristically defined loss that combines MAE loss and Pearson correlation coefficient loss. The latter is defined as:

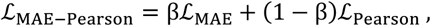

where ℒ_Pearson_ = 1 − *ρ* (*f*(***x***), ***y***) and *ρ* represents Pearson correlation coefficient. β is dynamically set as the constant value of *ρ* (*f*(***x***), ***y***) in each mini-batch iteration. Intuitively, when the Pearson correlation coefficient is high, the model focuses more on the MAE loss, and vice versa.

GARM was benchmarked compared to GAR also on four real world Perturb-seq datasets: HepG2^7^, Jurkat^7^, K562^6^, RPE1^6^. Similarly as synthetic experiments, we run 3 baselines of GAR with different method-specific hyper-parameter α in {0.1, 1, 10} as utilized in the original paper^14^, as oppose to running GARM without any method-specific hyper-parameter. Each dataset was randomly split with 75% of data used for training and the remaining 25% used only for testing. 10% of the training data is then selected randomly as the validation data. Each model was trained by Adam optimizer^17^ with initial learning rate tuned and set as 1e-3, running 30 total epochs with learning rate decreased by 10 folds at the end of 10-th and 20-th epoch. The batch size was fixed as 16 and weight decay tuned in {1e-4, 1e-6, 1e-8} per-(dataset, split). This procedure was repeated 5 times with independent random data splits and a different random seed. The reported best testing performance is chosen based on the best validation performance per-(dataset, split) for each compared methods with possible early stopping on training epochs and choice on weight decay. The backbone model was a 4-layered FFNN with 2 hidden layers for both GAR and GARM, where each layer contains 1024 neurons. ELU was used as the activation function^16^. GARM always maintains competitive performance with GAR tuned with 3 different hyper-parameter α; and for some cases, such as PrtR Spearman on Jurkat and MSE on K562, GARM enjoys better performance than all the variants of GAR (**Extended data Fig. 3a)**.

### Predicting the effect of unseen perturbations within a dataset

GARM was compared to four other methods that were developed to predict the results of Perturb-seq screens: GEARS^11^, scGPT^12^, GenePert^13^, and a co-expression based linear model that we refer to as ‘Coexpress’^15^.

All methods were tested for predicting the effects of unseen perturbations when trained and tested on the data obtained in the same Perturb-seq screen.

Four CRISPRi Perturb-seq datasets were used, performed in HepG2^7^, Jurkat^7^, K562^6^, and RPE1^6^ cells, with the latter two datasets also used in their curated form, where only a subset of strong perturbations and high-varying genes were retained^11^. HepG2 are epithelial-like cells isolated from a hepatocellular carcinoma of a 15-year-old white male; Jurkat are T cells isolated from the peripheral blood of a 14-year-old, male, acute T-cell leukemia patient; K562 are lymphoblast cells isolated from the bone marrow of a 53-year-old, female, chronic myelogenous leukemia patient; RPE1 are retinal pigment epithelial cell line isolated from a 1-year-old female.

In all datasets, the response to perturbation *p* was defined as

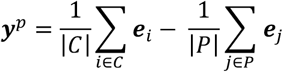

Where ***e*** is the log1p-transformed transcript-per-million (TPM) values, *C* denotes the control cells carrying a non-targeting control sgRNA, and *P* denotes all the cells with an sgRNA targeting the gene perturbed in perturbation *p*.

Each dataset was randomly split with 75% of data used for training and the remaining 25% used only for testing. 10% of the training data was then selected randomly as the validation data. For iterative methods, GEARS and GARM, the Adam optimizer^17^ was used for training.

The setup details of the compared methods are as following:

GEARS^11^: the initial learning rate was tuned and fixed as 1e-3 and the learning rate was decreased by half at the end of each training epoch (one-pass of the whole training data). The hidden dimension was tuned and fixed as 128. The total training epoch number was set as 10. Because GEARS was trained at single cell level and the datasets usually contains hundreds of single cells per perturbation, training GEARS requires hundreds more iterations than the method trained in pseudo-bulk level per epoch (**Supplementary, Practical running time analysis**). The training batch size was set as 32 and weight decay was tuned in {1e-4, 1e-6, 1e-8}. Because it is an iterative method, the evaluation was taken at early stopping epochs. The evaluation is made every 1 epoch, and early stopping is applied for the best validation performance.

Coexpress^15^: the hidden dimension for PCA (principal component analysis) was tuned in {32, 64, 128} and the L2-regularization for the linear regression model was tuned in {1e4, 1e3, 1e2, 1e1, 1e0, 1e-1, 1e-2, 1e-3, 1e-4, 1e-5, 1e-6}.

scGPT^12^: the nonlinear embedding for each genetic perturbation extracted from scGPT model was taken as the input for a ridge regression model for the output prediction, which showed better performance than the original perturbation prediction design via the scGPT model^13^. The L2-regularization for the ridge regression model was tuned in {1e4, 1e3, 1e2, 1e1, 1e0, 1e-1, 1e-2, 1e-3, 1e-4, 1e-5, 1e-6}.

GenePert^13^: the nonlinear embedding for each genetic perturbation extracted from GenePT model^18^ was taken as the input for a ridge regression model for the output prediction. The L2-regularization for the ridge regression model was tuned in {1e4, 1e3, 1e2, 1e1, 1e0, 1e-1, 1e-2, 1e-3, 1e-4, 1e-5, 1e-6}.

GARM: the nonlinear embedding extracted from GenePT model and PCA embedding extracted from GO for each genetic perturbation were taken as the input for a Feed-Forward Neural Network (FFNN) based backbone model (**Fig. 2a, Methods, Input Features for GARM**), where the layer number was set as 2 for both encoder and decoder with 1024 neurons in each layer. The initial learning rate was tuned and fixed as 1e-3 and the learning rate was decreased by 10-fold at the end of the 100^th^ and 200^th^ training epoch. The total training epoch number was set as 300 and the batch size was set as 16. The evaluation is made every 10 epochs, and early stopping is applied for the best validation performance.

For each dataset, the data was split independently 5 times with random seed in {1, 2, 3, 4, 5}; each method was run with its tuning hyper-parameters on each data split. The reported best testing performance was selected based on the best validation performance per (dataset, split) for each method with the choices on its hyper-parameters (and early stopping epochs in the case of iterative methods). For each method performances were quantified based on all data splits on each dataset (**Fig. 2b, Extended Data Fig. 3b**). The p-values are computed based on the one-side t-test for the 5 performance values from the 5 data splits.

For the detailed performance analysis on each gene (**Fig 3a, c, Extended Data Fig. 3c**), the best hyper-parameters were chosen for each method, namely, the highest average PrtR Pearson correlation coefficients across all genes in the validation dataset. GEARS was applied with a 1e-3 learning rate and 1e-6 weight decay; Coexpress was applied with 32 PCs (principal components) and 0.1 weight decay; scGPT was applied with 10.0 weight decay; GenePert was applied with a 1.0 weight decay; GARM was applied with 1e-3 learning rate and 1e-8 weight decay. The other settings are the same as described previously. The performance for each gene is averaged by the five data splits with different random seeds.

### Cross-dataset prediction of perturbation effects in an unseen cellular context

Four large CRISPRi Perturb-seq datasets were used for the cross-dataset predictions: HepG2^7^, Jurkat^7^, K562^6^, and RPE1^6^. Only 6,641 genes included in all 4 datasets were considered, using log_2_1p-transformed TPM values. In each iteration, one of the four datasets was used for testing, one for validation, and the remaining two datasets for training, resulting in 4 × 3 = 12 different data splits. When one dataset was used for testing, the evaluations were made from 3 different data splits by iterating the remaining 3 dataset as validation.

The methods were applied with the settings described above for the within-dataset experiments, except GEARS that was trained here at the pseudo-bulk level (due to high computational and memory cost on multiple datasets). The total training epoch number for GEARS was set here as 500 to make sure it is well-trained, where the learning rate was decreased by 0.997 at the end of each epoch and evaluation frequency is set as every 50 epochs with potential early stopping.

For the detailed performance analysis at the gene level (**Fig 3b, d, Extended Data Fig. 3e**) the best hyper-parameter was selected based on the average PrtR Pearson correlation coefficients across all genes. Weight decay of 1e-4, 0.1, 10.0, 1.0, and 1e-6 were used for GEARS, Coexpress, scGPT, GenePert and GARM, respectively. A learning rate of 1e-3 was used for GARM and GEARS, and 128 PCs were used in Coexpress. The other settings were the same as described previously. The performance for each gene was averaged across the three data splits.

### Simple and noninformative model

A baseline model that always predicts the perturbation response as the mean of all the perturbation responses from the training data^19^ was used here to evaluate prediction performances metrics. For cross-dataset experiment setting whose training part includes multiple different datasets, the mean is taken from all of them. We refer to this model as ‘PertMean’. For PrtR evaluations, PertMean is generally not evaluable because it always predicts a constant value per gene, which make the Pearson or Spearman correlation coefficients undefined. In that case, a uniformly random prediction between [0,1) is generated to provide a general bottom line for comparison.

### Input features for GARM

GARM utilizes external feature from GO (Gene Ontology) and ChatGPT (refined by GenePT^18^) for the feature of perturbation, ***x***^*p*^, following previous work^11,13^. The representations for gene from GenePT were already in vector form, which were directly fed into the FFNN based encoder. For GO data in graph format, where different gene terms are associated with different GO terms, PCA was applied to extract the low dimension (PCs = 256) representation vector for each gene. The two types of vectors were use as the input feature of different perturbations for GARM. Perturbations not represented in GO (15 out of 2057 or 20 out of 2393) were excluded throughout because GEARS relies on this input. Perturbations GenePT features were imputed based on the averaged value of the training perturbations. The GO terms associated to each gene were downloaded from https://dataverse.harvard.edu/api/access/datafile/6153417. GenePT features were downloaded from https://zenodo.org/records/10833191.

### Gene Set Enrichment Analyses (GSEA)

Gene Ontology (GO)^20,21^, REACTOME^22^, KEGG^23^, and other annotated gene sets from the Molecular Signature Database (MSigDB)^24^ were used. GSEAs were performed using the R Package fgsea (version 1.24.0) for each of the 4 datasets, with the genes ranked based on the within-dataset PrtR Pearson correlation values (**Fig. 4a, Supplementary Table 3**). As there were no gene sets identified as showing a significant skew to low PrtR values based on GSEA, another analysis was performed where genes with low PrtR values (< 0.1 Pearson correlation coefficient) in at least 2 out of the 4 datasets were identified as low PrtR genes. Hypergeometric tests were performed to examine which gene sets are enriched for low PrtR genes (**Supplementary Table 3**).

## DATA AND CODE AVAILABILITY

The HepG2^7^ and Jurkat^7^ Perturb-seq datasets were downloaded from the Gene Expression Omnibus (GEO; GSE264667). K562 and RPE1 Perturb-seq datasets were downloaded from [Figshare](https://doi.org/10.25452/figshare.plus.20029387.v1). GenePT features were downloaded from https://zenodo.org/records/10833191. The curated K562^6^ and RPE1^6^ Perturb-seq datasets were adopted from the GEARS paper, which is available on [Dataverse](https://doi.org/10.7910/DVN/BD93JY). The code developed here with ancillary files and instructions is available at https://github.com/Jerby-Lab/GARM.

## EXTENDED DATA FIGURES

**Extended Data Figure 1.**
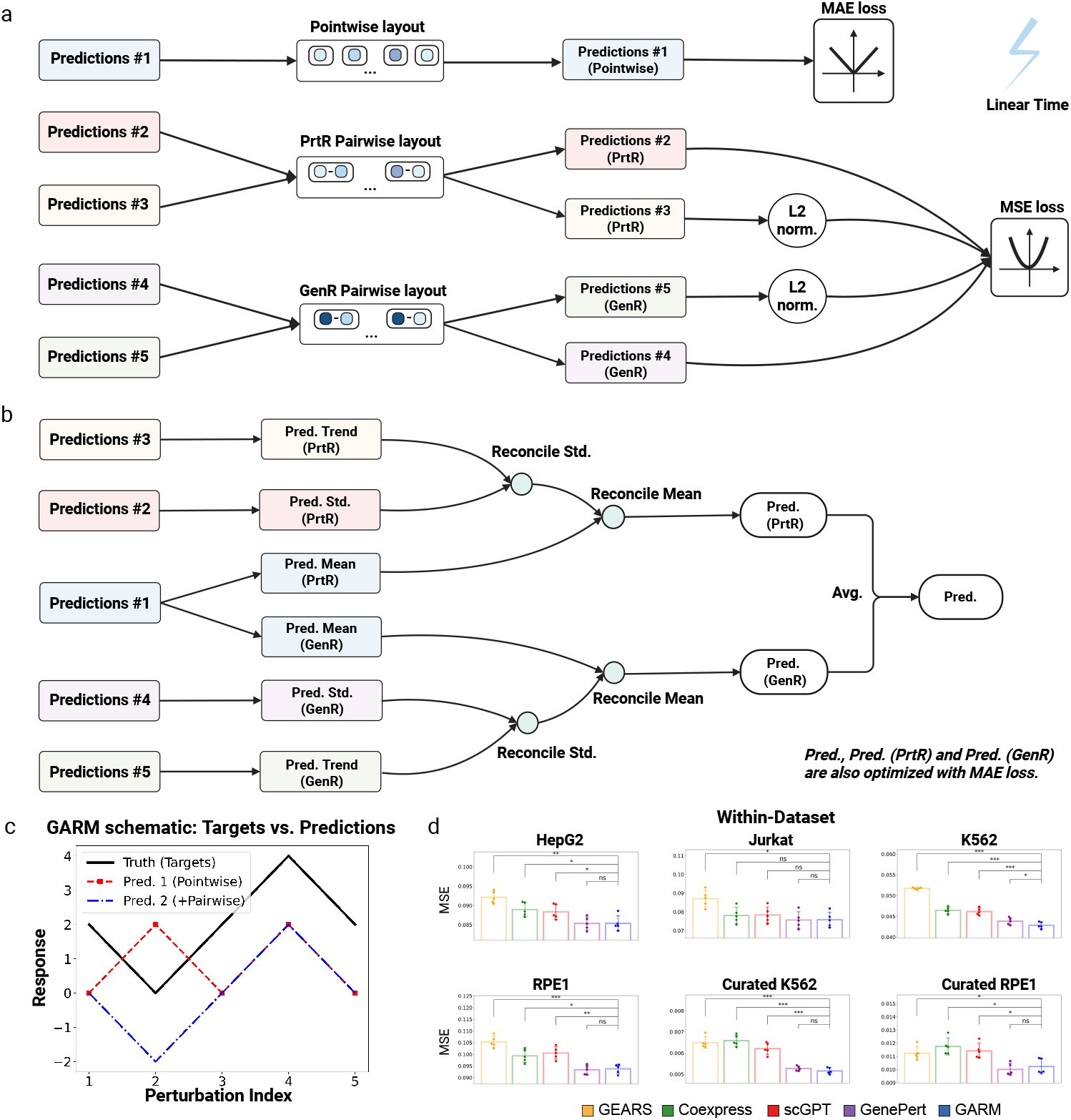
GARM design and prediction consolidation. **(a)** Design for GARM on the loss functions for different versions of the prediction. **(b)** Diagram for GARM on reconciling different predictions. **(c)** A graphical demonstration showing the advantage of pairwise loss over pointwise loss in model convergence. Under pointwise loss, such as MSE or MAE, predictions 1 and 2 have equal loss. However, under pairwise loss, prediction 2 has a smaller loss because the pairwise relationships are maintained. In this case, prediction 2 can then be shifted to a perfect fit with a simple bias-correction. **(d)** Comparison of different methods in the Within-Dataset setting for MSE evaluation (mean ± SD), showing the 5 performance values from the 5 data splits on different random seeds. ^*\*\*\**^*P* < 0.0001, ^**^*P* < 0.001, ^*^*P* < 0.05, one-sided t-test. GARM with pairwise loss demonstrates significantly better performance on K562 over all other competitors with pointwise evaluation, which might be benefited from the case in (c).

**Extended Data Figure 2.**
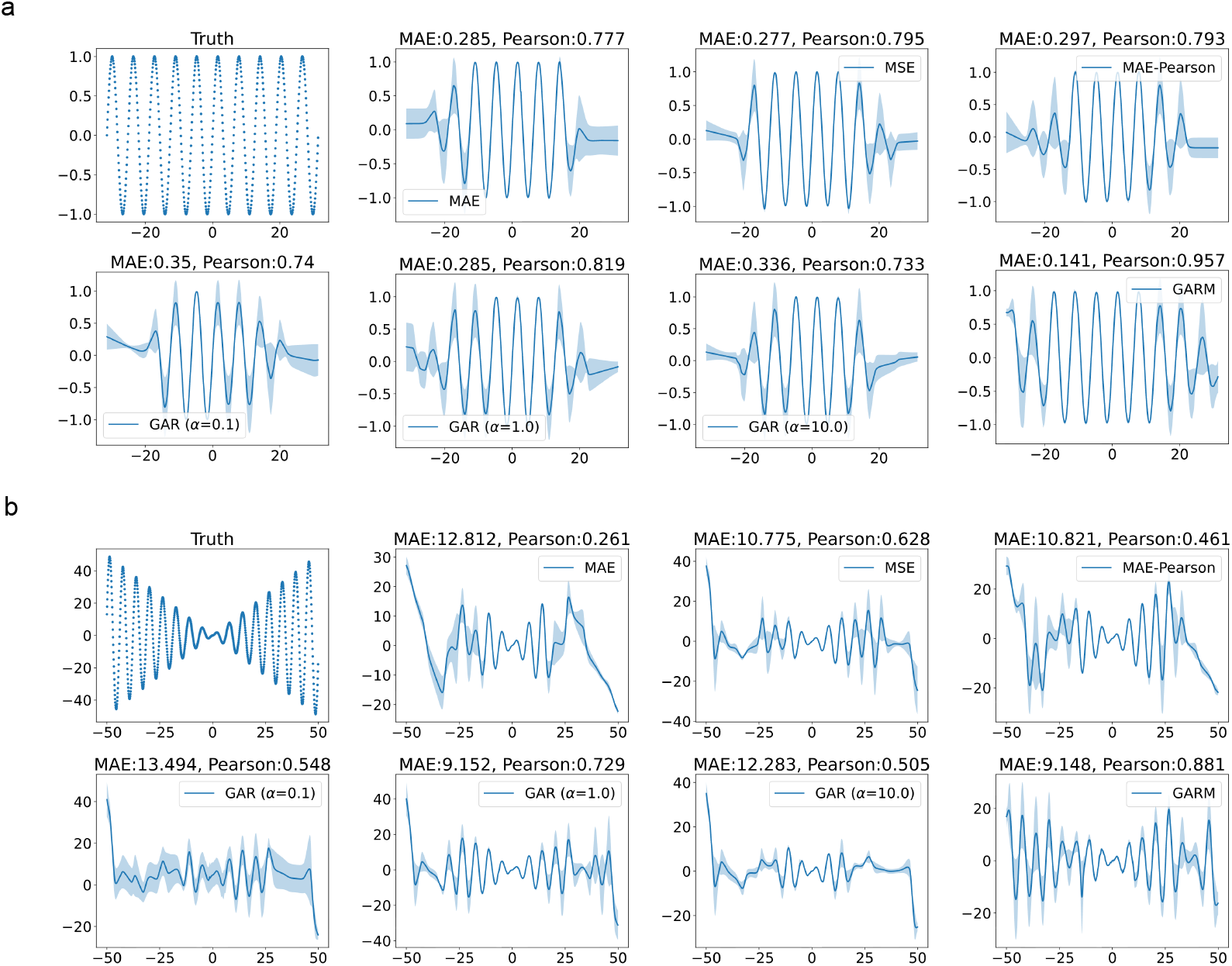
Benchmarking GARM using synthetic data. **(a)** Comparison between GARM vs. other regression methods on the Sine dataset. **(b)** Comparison between GARM vs. other regression methods on the Amplified-Sine dataset. For both (a) and (b), the second best is GAR, which requires hyperparameter tuning for α.

**Extended Data Figure 3.**
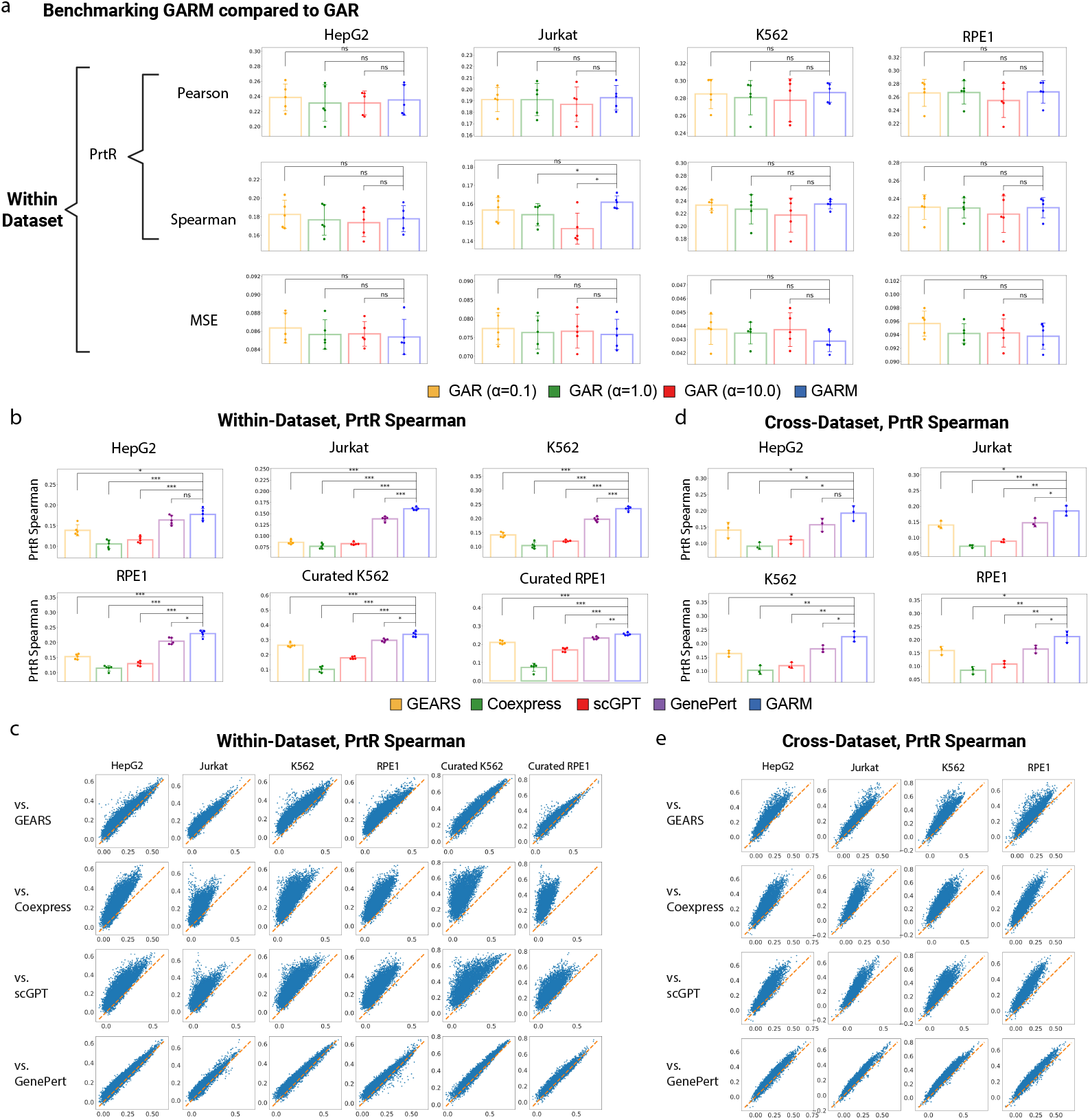
GARM benchmarked compared to GAR and other methods based on PrtR Spearman. **(a)** Comparison for GARM vs. GAR (with 3 different choices on hyper-parameter α) based on PrtR Pearson and Spearman correlation and MSE evaluations in Within-Dataset setting (mean ± SD). **(b-c)** Comparison of different methods in the Within-Dataset setting using PrtR Spearman correlation coefficients evaluation (mean ± SD), showing **(b)** 5 performance values from the 5 data splits on different random seeds, and **(c)** per gene performances of GARM (y axis) vs. the other method (x axis). **(d-e)** Comparison of different methods in the Cross-Dataset setting using PrtR Spearman correlation coefficients evaluation, showing **(d)** the 3 performance values from the 3 data splits, and **(e)** per gene performances of GARM (y axis) vs. the other method (x axis). For all barplots, ^***^*P* < 0.0001, ^**^*P* < 0.001, ^*^*P* < 0.05, one-sided t-test.

## SUPPELMENTARY TABLES

**Supplementary Table 1.** Within-dataset AUROC and AUPRC prediction evaluations. Average AUROC and AUPRC values obtained when predicting perturbations that up- and down-regulate each gene in the Within-dataset setting. The reported values are in mean (standard deviation) format across the 5 averaged values from the 5 different random splits of the data.

| Testing Performance |  | Average PrtR |  |  |  |
| --- | --- | --- | --- | --- | --- |
|  |  | AUROC (up) | AUPRC (up) | AUROC (down) | AUPRC (down) |
| Hepg2 | PertMean | 0.5 (0.0) | 0.2003 (0.0) | 0.5 (0.0) | 0.2003 (0.0) |
|  | GEARS | 0.5703 (0.005) | 0.2755 (0.0057) | 0.5721 (0.0063) | 0.2792 (0.0068) |
|  | Coexpress | 0.5577 (0.0045) | 0.2601 (0.0065) | 0.5555 (0.0038) | 0.2479 (0.0032) |
|  | scGPT | 0.5608 (0.0028) | 0.2656 (0.0034) | 0.559 (0.0047) | 0.2471 (0.0032) |
|  | GenePert | 0.5854 (0.006) | 0.2876 (0.0068) | 0.5852 (0.005) | 0.2821 (0.0067) |
|  | GARM | **0.5927 (0.0059)** | **0.3 (0.0078)** | **0.591 (0.0075)** | **0.2963 (0.0111)** |
| Jurkat | PertMean | 0.5 (0.0) | 0.2003 (0.0) | 0.5 (0.0) | 0.2003 (0.0) |
|  | GEARS | 0.5391 (0.0009) | 0.239 (0.0007) | 0.5476 (0.0028) | 0.2548 (0.0033) |
|  | Coexpress | 0.5457 (0.0015) | 0.2446 (0.0021) | 0.5359 (0.0047) | 0.2351 (0.0017) |
|  | scGPT | 0.5433 (0.0026) | 0.241 (0.0014) | 0.5412 (0.004) | 0.2342 (0.0034) |
|  | GenePert | 0.5665 (0.0021) | 0.2569 (0.0015) | 0.577 (0.0032) | 0.28 (0.0036) |
|  | GARM | **0.5766 (0.0052)** | **0.2738 (0.0065)** | **0.5932 (0.0038)** | **0.3009 (0.0073)** |
| K562 | PertMean | 0.5 (0.0) | 0.1996 (0.0) | 0.5 (0.0) | 0.1996 (0.0) |
|  | GEARS | 0.5751 (0.006) | 0.2821 (0.0094) | 0.5748 (0.0043) | 0.2813 (0.0044) |
|  | Coexpress | 0.5609 (0.0052) | 0.2695 (0.0056) | 0.5508 (0.0059) | 0.2387 (0.0049) |
|  | scGPT | 0.5678 (0.003) | 0.2771 (0.0061) | 0.5588 (0.0019) | 0.2407 (0.0008) |
|  | GenePert | 0.6042 (0.0045) | 0.3105 (0.0074) | 0.6046 (0.0043) | 0.3022 (0.005) |
|  | GARM | **0.6197 (0.0051)** | **0.3324 (0.0129)** | **0.6242 (0.0029)** | **0.3312 (0.0078)** |
| RPE1 | PertMean | 0.5 (0.0) | 0.2003 (0.0) | 0.5 (0.0) | 0.2003 (0.0) |
|  | GEARS | 0.5764 (0.004) | 0.2722 (0.0058) | 0.5815 (0.0055) | 0.2815 (0.0064) |
|  | Coexpress | 0.5547 (0.0045) | 0.2499 (0.004) | 0.558 (0.0061) | 0.2524 (0.0041) |
|  | scGPT | 0.5642 (0.004) | 0.2516 (0.0027) | 0.5656 (0.003) | 0.252 (0.0025) |
|  | GenePert | 0.6032 (0.0066) | 0.2929 (0.0062) | 0.6042 (0.0051) | 0.2957 (0.0055) |
|  | GARM | **0.6184 (0.0055)** | **0.3132 (0.0098)** | **0.6188 (0.0033)** | **0.3176 (0.0042)** |
| Curated K562 | PertMean | 0.4999 (0.0) | 0.2022 (0.0) | 0.5 (0.0001) | 0.2023 (0.0) |
|  | GEARS | 0.6566 (0.0079) | 0.3805 (0.0103) | 0.6192 (0.0052) | 0.3201 (0.0053) |
|  | Coexpress | 0.5436 (0.0064) | 0.2607 (0.0104) | 0.5288 (0.0014) | 0.233 (0.001) |
|  | scGPT | 0.6114 (0.005) | 0.3281 (0.0086) | 0.5748 (0.0045) | 0.2631 (0.0038) |
|  | GenePert | 0.672 (0.0061) | 0.4027 (0.0087) | 0.637 (0.0045) | 0.3296 (0.0052) |
|  | GARM | **0.6911 (0.0084)** | **0.428 (0.0117)** | **0.6591 (0.0049)** | **0.362 (0.0055)** |
| Curated RPE1 | PertMean | 0.5 (0.0001) | 0.2006 (0.0) | 0.5 (0.0) | 0.2005 (0.0) |
|  | GEARS | 0.6151 (0.0063) | 0.3228 (0.0059) | 0.601 (0.006) | 0.3077 (0.0061) |
|  | Coexpress | 0.5364 (0.0056) | 0.2352 (0.0053) | 0.5344 (0.0009) | 0.2356 (0.004) |
|  | scGPT | 0.5898 (0.006) | 0.2833 (0.0049) | 0.583 (0.004) | 0.2736 (0.0026) |
|  | GenePert | 0.6275 (0.0025) | 0.3294 (0.0051) | 0.6154 (0.0057) | 0.3113 (0.0067) |
|  | GARM | **0.6359 (0.004)** | **0.3452 (0.0032)** | **0.6279 (0.0044)** | **0.3315 (0.0061)** |

**Supplementary Table 2.**
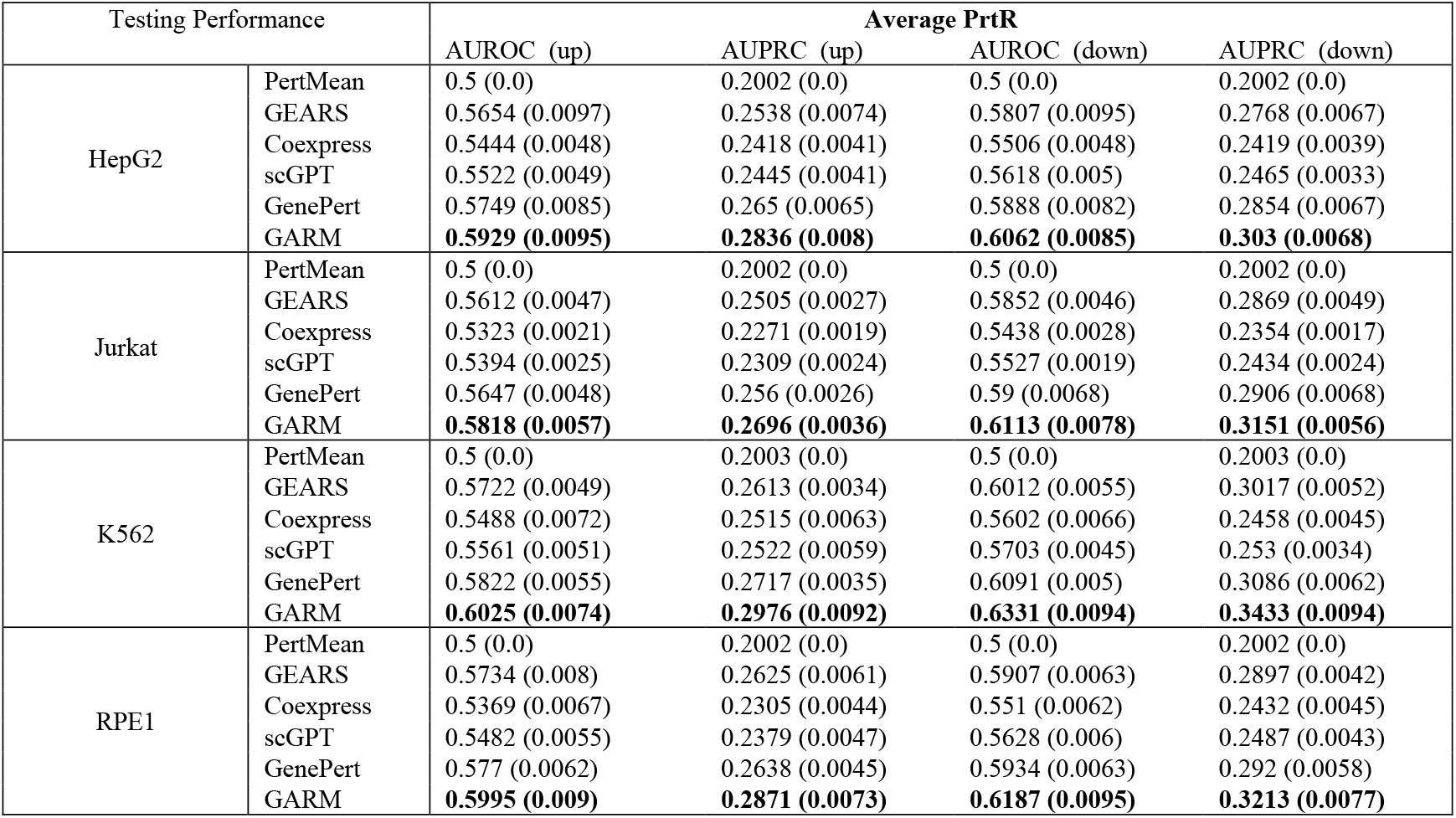
Cross-dataset AUROC and AUPRC prediction evaluations. Average AUROC and AUPRC values obtained when predicting perturbations that up- and down-regulate each gene in the cross-dataset setting. The reported values are in mean (standard deviation) format across the 3 averaged values from the 3 different data training and validation splits.

| Testing Performance |  | Average PrtR |  |  |  |
| --- | --- | --- | --- | --- | --- |
|  |  | AUROC (up) | AUPRC (up) | AUROC (down) | AUPRC (down) |
| HepG2 | PertMean | 0.5 (0.0) | 0.2002 (0.0) | 0.5 (0.0) | 0.2002 (0.0) |
|  | GEARS | 0.5654 (0.0097) | 0.2538 (0.0074) | 0.5807 (0.0095) | 0.2768 (0.0067) |
|  | Coexpress | 0.5444 (0.0048) | 0.2418 (0.0041) | 0.5506 (0.0048) | 0.2419 (0.0039) |
|  | scGPT | 0.5522 (0.0049) | 0.2445 (0.0041) | 0.5618 (0.005) | 0.2465 (0.0033) |
|  | GenePert | 0.5749 (0.0085) | 0.265 (0.0065) | 0.5888 (0.0082) | 0.2854 (0.0067) |
|  | GARM | **0.5929 (0.0095)** | **0.2836 (0.008)** | **0.6062 (0.0085)** | **0.303 (0.0068)** |
| Jurkat | PertMean | 0.5 (0.0) | 0.2002 (0.0) | 0.5 (0.0) | 0.2002 (0.0) |
|  | GEARS | 0.5612 (0.0047) | 0.2505 (0.0027) | 0.5852 (0.0046) | 0.2869 (0.0049) |
|  | Coexpress | 0.5323 (0.0021) | 0.2271 (0.0019) | 0.5438 (0.0028) | 0.2354 (0.0017) |
|  | scGPT | 0.5394 (0.0025) | 0.2309 (0.0024) | 0.5527 (0.0019) | 0.2434 (0.0024) |
|  | GenePert | 0.5647 (0.0048) | 0.256 (0.0026) | 0.59 (0.0068) | 0.2906 (0.0068) |
|  | GARM | **0.5818 (0.0057)** | **0.2696 (0.0036)** | **0.6113 (0.0078)** | **0.3151 (0.0056)** |
| K562 | PertMean | 0.5 (0.0) | 0.2003 (0.0) | 0.5 (0.0) | 0.2003 (0.0) |
|  | GEARS | 0.5722 (0.0049) | 0.2613 (0.0034) | 0.6012 (0.0055) | 0.3017 (0.0052) |
|  | Coexpress | 0.5488 (0.0072) | 0.2515 (0.0063) | 0.5602 (0.0066) | 0.2458 (0.0045) |
|  | scGPT | 0.5561 (0.0051) | 0.2522 (0.0059) | 0.5703 (0.0045) | 0.253 (0.0034) |
|  | GenePert | 0.5822 (0.0055) | 0.2717 (0.0035) | 0.6091 (0.005) | 0.3086 (0.0062) |
|  | GARM | **0.6025 (0.0074)** | **0.2976 (0.0092)** | **0.6331 (0.0094)** | **0.3433 (0.0094)** |
| RPE1 | PertMean | 0.5 (0.0) | 0.2002 (0.0) | 0.5 (0.0) | 0.2002 (0.0) |
|  | GEARS | 0.5734 (0.008) | 0.2625 (0.0061) | 0.5907 (0.0063) | 0.2897 (0.0042) |
|  | Coexpress | 0.5369 (0.0067) | 0.2305 (0.0044) | 0.551 (0.0062) | 0.2432 (0.0045) |
|  | scGPT | 0.5482 (0.0055) | 0.2379 (0.0047) | 0.5628 (0.006) | 0.2487 (0.0043) |
|  | GenePert | 0.577 (0.0062) | 0.2638 (0.0045) | 0.5934 (0.0063) | 0.292 (0.0058) |
|  | GARM | **0.5995 (0.009)** | **0.2871 (0.0073)** | **0.6187 (0.0095)** | **0.3213 (0.0077)** |

**\*Supplementary Table 3. Gene sets enriched for easy and hard to predict genes. (a-d)** GSEA of genes ranked based on PrtR Pearson correlation coefficients of GARM applied to (a) HepG2, (b) Jurkat, (c) K562, and (d) RPE1 Perturb-seq dataset. (e) GSEA (hypergeometric enrichment) for genes with low (<0.1 PrtR Pearson) in at least 2 out of the 4 datasets.

*Provided in a separate excel file.

## SUPPLEMENTARY FIGURES

## SUPPLEMENTARTY INFORMATION

### Practical running time analysis

As explained previously, the pairwise regression losses utilized in this work only require linear time computation complexity, which is the same as pointwise regression losses, such as MAE and MSE losses. In this subsection, we practically demonstrate the running time cost for GARM vs. other methods. We conduct the experiments on a server node with AMD EPYC 7713 64-Core Processor, CPU MHz: 1500.00, BogoMIPS: 3999.93, NVIDIA A100 80GB PCIe GPU card. For iterative methods, GEARS and GARM, we run 10 epochs for GEARS at single cell level and 300 epochs for GARM at pseudo-bulk level, which are consistent with our main experiment setting. The running time are summarized per epoch and for a full training process. For analytical methods, scGPT, GenePert and Coexpress, we directly solve the problem with matrix operations, such as matrix inverse and multiplication. We repeat the analytical methods with 10 different random seeds; then report the means and standard deviations. The HepG2^7^ dataset is utilized for analyzing the practical running time.

For GEARS, it takes 621.575 (104.846) seconds per epoch, which takes 6215.746 seconds in total for a full training process. For Coexpression, it takes 23.572 (1.715) seconds for a full analytical process. For scGPT and GenePert, they take 0.009 (0.0) and 0.059 (0.003) for a full analytical process. For GARM, it takes 1.127 (0.063) seconds per epoch, which takes 338.002 seconds in total for a full training process.

GEARS is proposed at single cell level, which takes much longer time than the other methods. Coexpress requires extra PCA computation cost, which is the main reason that it takes longer time than other analytical methods, i.e. scGPT and GenePert. It is also worth noting that we can potentially accelerate Coexpress PCA computation by utilizing GPU resource with alternative computation library. For the current implementation, we follow the original paper to use eigsh function scipy.sparse.linalg library.

